# Investigating the genetic and environmental basis of head micromovements during MRI

**DOI:** 10.1101/2021.10.25.465703

**Authors:** Frauke Beyer, Katrin Horn, S. Frenzel, Edith Hofer, Maria J. Knol, Filip Morys, Uku Vainik, Sandra Van der Auwera, Katharina Wittfeld, Yasaman Saba, Hieab HH Adams, Robin Bülow, Hans Grabe, Georg Homuth, Marisa Koini, Markus Loeffler, Helena Schmidt, Reinhold Schmidt, Alexander Teumer, MW Vernooij, Arno Villringer, Henry Völzke, Hazel Zonneveld, A. Dagher, Markus Scholz, AV Witte, on behalf of the NeuroCHARGE working group

**Author notes:** Contribution Study conception, Data analysis, Manuscript writing. Contribution genomic data and MetaGWAS analysis. Contribution phenotype preparation, statistical analysis. Contribution bioinformatics, statistical analysis. Contribution phenotype data aquisition, funding. Contribution MRI data aquisition. Contribution genotype data aquisition. Contribution phenotype preparation. Contribution study conception, funding. Contribution genotype data acquisition, funding. Contribution genotype data preparation. Contribution phenotype data aquisition, data processing. Contribution Study conception, Discussion and revision, funding. Contribution genotype data acquisition, data analysis, study conception, funding. Contribution Study conception, Manuscript writing and revision, Discussion. <u>Data availability statement:</u> Summary statistics for the GWAS meta-analysis on head motion will be deposited in the Leipzig Health Atlas (https://www.health-atlas.de/). Researchers may obtain access to the raw data for UKBB, LIFE-Adult and the Rotterdam study via data usage requests, for all other studies the authors may be contacted to provide data access. <u>Funding statements</u>. **LIFE-Adult** The LIFE-Adult was funded by the Leipzig Research Center for Civilization Diseases (LIFE). LIFE is an organizational unit affiliated to the Medical Faculty of the University of Leipzig. LIFE is funded by means of the European Union, by the European Regional Development Fund (ERDF) and by funds of the Free State of Saxony within the framework of the excellence initiative (Grant Numbers: 713-241202, 713-241202, 14505/2470, 14575/2470). Analysis was also funded by the Deutsche Forschungsgemeinschaft (Grant Number: CRC 1052 “Obesity mechanisms” A1). **UKBB** The UK Biobank and its Imaging Enhancement are funded by the Medical Research Council and the Wellcome Trust. Data was accessed through agreement 35605 (PI, A. Dagher). Uku Vainik has been funded by Estonian Research Council’s personal research funding start-up grant PSG656. <u>Ethics approval statement</u> All included studies were performed according to the declaration of Helsinki and received approval of local ethic committees.

## Abstract

**Introduction:** Head motion during magnetic resonance imaging is heritable. Further, it shares phenotypical and genetic variance with body mass index (BMI) and impulsivity. Yet, to what extent this trait is related to single genetic variants and physiological or behavioral features is unknown. We investigated the genetic basis of head motion in a meta-analysis of genome-wide association studies. Further, we tested whether physiological or psychological measures, such as respiratory rate or impulsivity, mediated the relationship between BMI and head motion.

**Methods:** We conducted a genome-wide association meta-analysis for mean and maximal framewise head displacement (FD) in seven population neuroimaging cohorts (UK Biobank, LIFE-Adult, Rotterdam Study cohort 1-3, Austrian Stroke Prevention Family Study, Study of Health in Pomerania; total N = 35.109). We performed a pre-registered analysis to test whether respiratory rate, respiratory volume, self-reported impulsivity and heart rate mediated the relationship between BMI and mean FD in LIFE-Adult.

**Results:** No variant reached genome-wide significance for neither mean nor maximal FD. Neither physiological nor psychological measures mediated the relationship between BMI and head motion.

**Conclusion:** Based on these findings from a large meta-GWAS and pre-registered follow-up study, we conclude that the previously reported genetic correlation between BMI and head motion relies on polygenic variation, and that neither psychological nor simple physiological parameters explain a substantial amount of variance in the association of BMI and head motion. Future imaging studies should thus rigorously control for head motion at acquisition and during preprocessing.

## 1. Introduction

Head motion (HM) during magnetic resonance imaging (MRI) is a major source of nuisance in MRI studies as it decreases data quality and induces significant bias in imaging biomarkers such as functional connectivity and gray matter volume (Power et al., 2012; Savalia et al., 2017; Van Dijk et al., 2012). There is a remarkably consistent association of higher body mass index (BMI) and higher head motion, yet the underlying mechanisms remain poorly understood (Beyer et al., 2017; Hodgson et al., 2016; Madan, 2018).

Previous studies have attributed the association to physiological parameters such as breathing rate and chest volume (Ekhtiari et al., 2019; Siegel et al., 2017). Higher respiratory rate, shallower breaths and lower lung volume have been consistently reported in obesity (Burki & Baker, 1984; Littleton, 2012). Respiration explains a considerable part of variance in head motion and signal variation during functional MRI, which is on the one hand due to physical motion associated with breathing and on the other hand related to artificial head motion induced by chest motion and subtle susceptibility shifts (Fair et al., 2020). Thus, differences in respiratory parameters might mediate the observed BMI-head motion relationship.

A different line of research suggests a neurobiological trait, which predisposes to higher head motion and potentially more impulsive (motor) behavior in general (Zeng et al., 2014). Higher head motion has been consistently reported in attention-deficit disorder (ADHD) and is phenotypically and genetically correlated with impulsivity (Couvy-Duchesne et al., 2016; Kong et al., 2014; Thomson et al., 2020). Impulsivity and decreased inhibitory control show small yet reliable association with BMI, which could be in part explained by shared genetic factors (Meule & Blechert, 2016; Vainik et al., 2018). Head motion is moderately heritable (h^2^ ∼ 0.4) and shares genetic variance with BMI (ρ_g_ ∼ 0.8) in studies based on family structure (Couvy-Duchesne et al., 2014; Engelhardt et al., 2017; Hodgson et al., 2016). Thus, a genetically determined tendency towards impulsive behavior might underlie both BMI and head motion. Another psychological factor to be considered is anxiety and claustrophobic feelings which people with higher BMI might experience more often than lean participants in the narrow scanner bore. Anxiety (reflected by higher heart rate) might also contribute to higher head motion (van Minde et al., 2014).

Taken together, different psychological and physiological factors might contribute to the robust phenotypic association of BMI and head motion. Here, we aimed to further explore the role of these factors by

1. investigating the genetic basis of head motion in a meta-analysis of genome wide association studies (GWAS). We hypothesized to detect either body weight-related (e.g. FTO) or impulsivity/ADHD-related variants (e.g. ADGRL3), either one of which would be indicative of the pathways involved in the genetic association between head motion and BMI.
2. testing whether respiratory parameters, heart rate or measures of impulsivity mediated the association of BMI and head motion.

## 2. Methods

### 2.1 Meta-GWAS

We performed a GWAS meta-analysis of 35109 participants of European ancestry from 7 studies (stage 1) that contributed summary statistic data.

#### 2.1.1 Study populations

All participating studies except for the UK biobank study are part of the Cohorts for Heart and Aging Research in Genomic Epidemiology (CHARGE) consortium. Each study was approved by the local ethics committee and conducted according to the declaration of Helsinki. Informed consent was obtained from all participants.

The analysis plan for the meta-GWAS was pre-specified and sent to all participating studies (see https://osf.io/ahfv9/ for more details). Study participants were older than 18 years and not diagnosed with stroke, brain pathologies (e.g. tumors or traumatic brain injury) or dementia. For basic demographics of the participants see Supplementary Table 1.

#### 2.1.2 Imaging

Resting-state fMRI was performed on 1.5 or 3 Tesla scanners at all study sites. The phenotypes of interest mean and maximal frame-wise displacement (mFD, maxFD) were calculated according to Power et al. (Power et al., 2012). All but one study used FSL’s function MCFLIRT to extract 6 motion parameters (three rotational and three translational) which model the frame-to-frame head motion. In SHIP, AFNI’s 3dvolreg was used to calculate these parameters. Then, framewise displacement was calculated from these parameters according to (Power et al., 2012). We provided a publicly available custom script to all research groups (https://github.com/fBeyer89/life_followup_preproc/blob/master/qa/resting/qa_pipeline/utils.py). Information on scanner manufacturers, acquisition protocols and motion correction tools are provided in Supplementary Table 1.

#### 2.1.3 Genotyping and Imputation

Information on genotyping platforms, quality control procedures and imputations methods for each participating study are provided in Supplementary Table 2. All studies used commercially available genotyping arrays, including Illumina or Affymetrix arrays. Single study quality control was performed at the discretion of the single study groups. Using validated software, each study performed genotype imputations.

#### 2.1.4 Quality control of single study association results

Summary statistics of all studies were checked and harmonized with the software EasyQC. We discarded SNPs not in the reference panel (1000 Genomes phase 3, version 5, European ancestry), mismatching alleles or mismatching chromosomal position with respect to the reference. Moreover, we removed SNPs with missing allele information (effect allele, effect allele frequency), missing association statistics (beta estimates, standard errors) or missing imputation quality score. SNPs were filtered for weighted minor allele frequency (MAF) >1%. Genotyped SNPs were filtered for call rate >97% and p-value of Hardy-Weinberg test >10^−6^. Imputed SNPs were filtered for imputation quality score >0.5 and for deviation from reference allele frequency <20%. Finally, the alleles were harmonized so that the same effect allele was used in all studies. Variance inflation factor lambda was calculated for single study GWAS. Test statistics were corrected by genomic control if λ >1.

#### 2.1.5 Statistical Analysis

The single-study GWAS were performed locally following a uniform analysis plan provided to all study groups. We performed linear regression analysis of the log-transformed phenotypes of interest (mFD and maxFD). Covariates were age, sex, total intracranial volume (TIV) as well as three principle components of genetic population structure.

An additive gene-dose model was assumed. The analyses were performed for all individuals combined, and separately for female and male participants (self-reported; note that diverse gender was not assessed in most studies) following the SAGER guidelines (Heidari et al., 2016).

#### 2.1.6 Meta-analysis

Altogether two traits were meta-analysed, combined together or stratified by sex. Fixed effects inverse variance meta-analysis was performed as primary statistics. Random effects meta-analysis results were also reported. Meta-analysis results were filtered for number of contributing studies >2 and heterogeneity I^2^<0.75.

A p-value of <5 10^−8^ was considered genome-wide significant.

### 2.2 Investigation of respiratory and psychological measures in the relationship of BMI and head motion

We pre-registered this part of the analysis on the Open Science Framework .https://osf.io/rh52s.

We hypothesized that respiratory rate, respiratory volume per time (RVT), heart rate, total impulsivity, motor, or self-control impulsivity might mediate the link between BMI and HM. Further, we aimed to test whether BMI was associated with differences in the correlation between frame-to-frame HM and BOLD signal intensity with respiratory trace and RVT. Our sample was taken from the LIFE-Adult study, a population-based cohort study with 10.000 participants which included genotyping, MRI and deep phenotyping (Loeffler et al., 2015)(Engel et al., 2021 in preparation). The study was approved by the Ethics committee of the Medical Faculty of Leipzig University and all participants signed written informed consent and received a renumeration for their participation.

#### 2.2.1 Sample

We included 1006 participants with follow-up MRI acquisition available on June, 30th 2021. The flowchart in Supplementary Figure 8 illustrates the exclusion and missing data which led to the final sample sizes in the different analysis parts.

#### 2.2.2 Imaging and physiological data acquisition

Resting state fMRI was acquired with the same echo-planar-imaging sequence as in the baseline assessment (repetition time, 2 s; echo time, 30 ms; flip angle, 90°; image matrix, 64 × 64; 30 slices; field of view, 192 × 192 × 144 mm3, no multiband, iPAT acceleration: 1, voxel size of 3 mm × 3 mm, slice thickness of 4 mm, slice gap of 0.8 mm; 300 volumes; total acquisition time, 10:04 minutes). Physiological parameters were recorded during the scan using Siemens proprietary hardware (breathing belt and an oximeter attached to the index finger). The sampling rate was 50 Hz for both devices..

#### 2.2.3 rsfMRI and physiological data preprocessing

We used FSL’s function MCFLIRT to extract 6 motion parameters from the rsfMRI scan (three rotational and three translational) which model the frame-to-frame head motion. Framewise displacement was calculated from these parameters according to (Power et al., 2012) using a publicly available custom script (https://github.com/fBeyer89/life_followup_preproc/blob/master/qa/resting/qa_pipeline/utils.py). We used log10-transformed average mFD and frame-to-frame FD for analysis. We also calculated DVARS, a measure of BOLD fluctuations from frame to frame which allows us to quantify the effect of head motion/physiological parameters on the acquired signal (Power et al., 2015). Standardized DVARS was calculated from the raw fMRI scan with the ComputeDVARS function from the nipype.algorithms.confounds module in this script.

We used PhysIO toolbox (“Created: 2011-08-01”) in Matlab version 9.3 to preprocess the physiological data (Kasper et al., 2017). We performed automated pulse detection for detecting cardiac phase with default parameters for heartbeat duration outliers. Heart rate was estimated by averaging heart beat duration in a sliding window of 6 seconds (3 TR) (Chang et al., 2009). Respiration maximal and minimal amplitude (corresponding to respiratory cycles) was also automatically detected. Then, respiratory volume per time (RVT) was calculated by dividing the amplitude between adjacent breathing minima and maxima by the interpolated time interval between them (Birn et al., 2006). We normalized RVT by dividing it by the maximal RVT value for each participant and calculated the average and standard deviation of the normalized RVT. Respiratory rate (RR) was calculated as the inverse of the average of all interpolated breathing cycle durations. To estimate the correlation of head motion and respiration, we first downsampled the respiratory amplitude trace and RVT to the scan frequency (1/TR). Then, we calculated Pearson’s correlation of the respiratory trace/RVT with FD or DVARS. We excluded participants with physiologically implausible values of RR (< 0.16 Hz or > 0.5 Hz) and HR (< 40 bpm or > 110 bpm).

BMI was determined based on the anthropometric assessment in the LIFE-study and calculated as weight divided by height squared. We excluded participants with physiologically implausible values of BMI (<15 kg/m2 and >60 kg/m2).

#### 2.2.4 Psychological measures

We used the Barratt impulsiveness scale in its revised version (BIS-11) to assess self-reported impulsivity. Based on published scoring scheme, we derived the total impulsivity scale, the motor impulsivity subscale and the self-control subscale (Kong et al., 2014; Patton et al., 1995).

#### 2.2.5 Statistical analysis

All statistical analyses were done using R version 3.6.1. The code for the statistical analyses is publicly accessible at https://github.com/fBeyer89/Mediators_of_HeadMotion. As pre-registered, we performed mediation analyses for respiratory and psychological variables. BMI was the independent predictor, mFD was the outcome and all models were adjusted for age and sex (male as reference category). Mediating factors were RR, mean RVT and HR for the physiological and BIS_motor, BIS_selfcontrol, BIS_total for the psychological models.

The mediation models were implemented in the R package lavaan 0.6.7 and estimated using maximum likelihood. The standard errors and confidence intervals were derived from 1000 bootstraps. Depending on the number of tests per group, we used Bonferroni-corrected p-values (physiological: p<0.0167 and psychological mediations p<0.0167). Further, we tested whether the correlation of respiratory parameters and frame-to-frame head motion and BOLD signal variation was associated with BMI. To this end, we first derived the frame-to-frame correlation of downsampled RVT and the respiratory trace with frame-to-frame head motion and BOLD signal variation. Then, we estimated the association of BMI and these correlations in linear models adjusting for age and sex. We performed full-null model comparison (omitting BMI from the null models) and used a Bonferroni-adjusted p-value (p<0.0125) for the four models tested.

### Evaluation of statistical model assumptions

We checked all variables visually for normality and noticed no strong deviation except for BIS11_selfcontrol which was not normally distributed because only six items were used for its construction..

We visually inspected the residuals of the linear models used in the mediation analysis, and checked for variance inflation and heteroskedasticity. All residuals looked approximately normally distributed, and there was no case of strong heteroskedasticity or variance inflation with VIF >10.

We repeated the mediation models without influential cases defined by Cooks distance > 4/n (n is the number of individuals included in the respective models).

In the pre-registration we planned to check the quality of fit of the mediation models with common fit indices used in structural equation models (SEM). However, this was considered unnecessary since no latent variables were considered

## 2.3 Results

### 2.3.1 Meta-GWAS

Age and sex distribution as well as average mFD of the participating studies can be found in Supplementary Table 1.

Meta-analysis of seven studies and 35,109 subjects did not result in genome-wide significant associations neither for mFD nor max FD (see Figure 1). This applies for the combined and the stratified analyses (see Supplementary Figures 3 -- 6).

**Figure 1:**
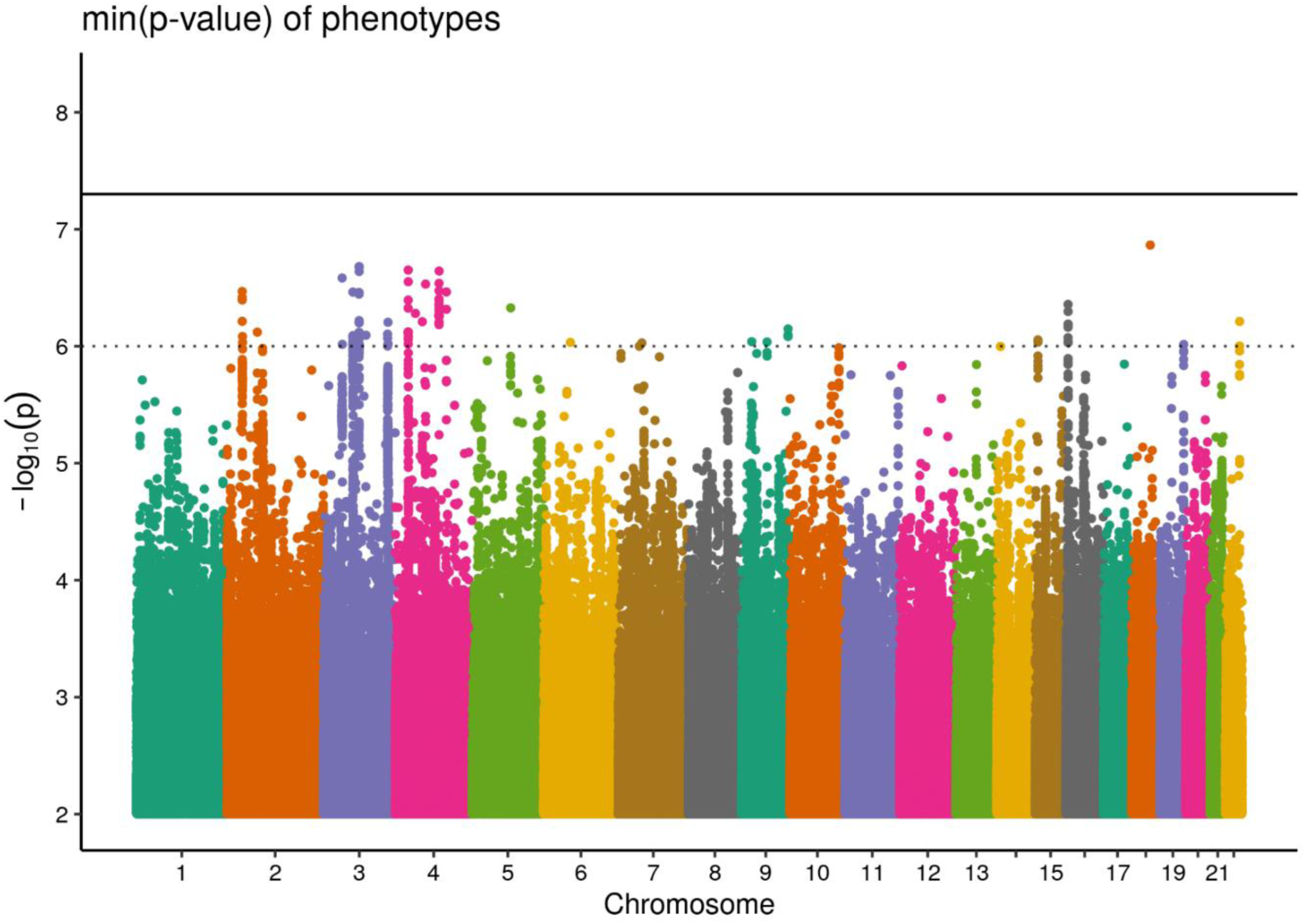
Manhattan plot showing the min(p-value) for an association of both phenotypes mean and maximal FD with genetic variants.

For mFD, there were 109 SPNs with a p<10^−6^. The strongest effects were on chromosome 3 in the variants rs2218592 (p=2.1 10^−7^) and rs9867092 (p=2.1 10^−7^) and on chromosome 16 in the variants rs8054401 (p=3.9 10^−7^) and rs741695 (p=5 10^− 7^) (see Supplementary Figures 1 and 2). According to the UK Biobank gene atlas (http://geneatlas.roslin.ed.ac.uk/phewas), the chr 3 variant is related to tea intake, while the chr 16 variant has strongest associations with body weight.

For maxFD, the only association found when thresholding for p<10^−6^ was on chromosome 4 with variants rs114571645 (p=2.3 10^− 7^) and rs6845933 (p=2.9 10^− 7^) (see Supplementary Figure 3). Variant rs114571645 is related to allergy according to the gene atlas.

In the analysis separated into males and females, also no variant reached genome-wide significance (see Supplementary Figures 5 – 8).

### 2.3.2 Mediation analyses

#### Physiological Mediation

There was no significant mediation effect of RR, RVT or HR on the relation between between BMI and mFD, also after excluding influential cases (see Table 1).

**Table 1:**
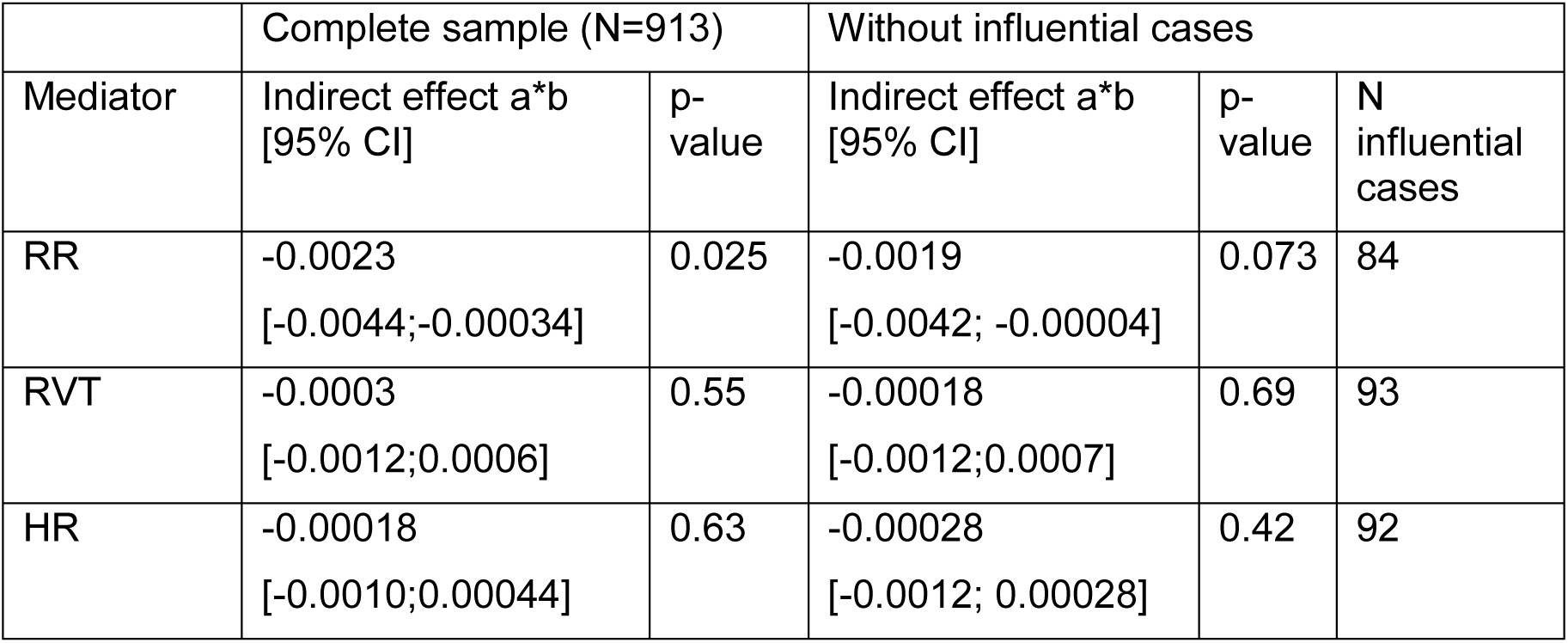
Results of the physiological mediation models, shown for the full sample and without influential cases. RR: respiratory rate; RVT: respiratory volume per time, HR: heart rate, CI: Confidence interval.

| Mediator | Complete sample (N=913) |  | Without influential cases |  |  |
| --- | --- | --- | --- | --- | --- |
|  | Indirect effect a*b<br>[95% CI] | p-value | Indirect effect a*b<br>[95% CI] | p-value | N influential cases |
| RR | -0.0023<br>[-0.0044;-0.00034] | 0.025 | -0.0019<br>[-0.0042; -0.00004] | 0.073 | 84 |
| RVT | -0.0003<br>[-0.0012;0.0006] | 0.55 | -0.00018<br>[-0.0012;0.0007] | 0.69 | 93 |
| HR | -0.00018<br>[-0.0010;0.00044] | 0.63 | -0.00028<br>[-0.0012; 0.00028] | 0.42 | 92 |

All mediation models showed a strong direct effect between BMI and mFD (for RR: unstandardized β=0.049, 95% confidence interval (CI): [0.043;0.054], p<0.001). BMI was significantly associated with higher HR (unstandardized β= 0.28, 95% CI: [0.12;0.46], p<0.001), but not RR (unstandardized β= 0.00092,, 95% CI: [0.00013; 0.0017], p=0.018) or RVT (unstandardized β=0.00076; confidence interval (CI): [-0.0018; 0.0031], p=0.54). Further, both respiratory parameters were strongly negatively associated with mFD (RR: unstandardized β=-2.49, 95% CI: [-3.0;-1.99], p<0.001; RVT: unstandardized β=-0.35, 95% CI: [-0.49;-0.21], p<0.001; see Figure 2).

**Figure 2:**
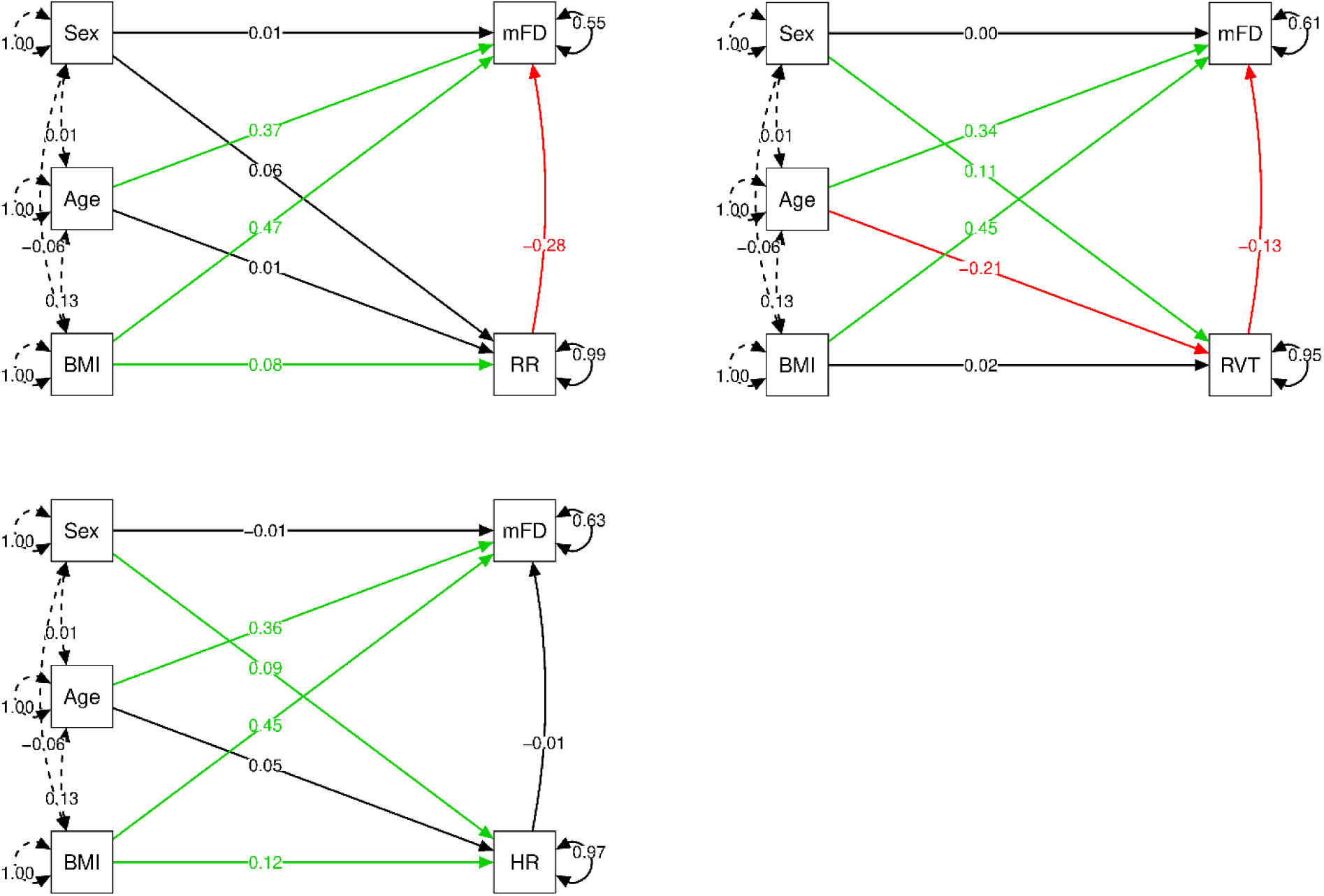
Mediation models for RR (upper left panel), RVT (upper right panel) and HR (lower panel) showing standardized estimates. Dashed arrows indicate residual (co-)variance. Solid arrows indicate estimated regression coefficients, green/red indicate significant positive/negative associations (p<0.05), black indicates no significant association. The reference category for sex was male. BMI: body mass index, mFD: mean framewise displacement, RR: respiratory rate, RVT: respiratory volume per time, HR: heart rate.

#### Regression analysis

There was a significant negative association of higher BMI and the correlation between framewise FD and RVT (β=-0.0058, 95% C.I.: [-0.0088;-0.0027], p=0.00025) which remained significant after excluding 43 influential cases. There was no significant association with any of the other framewise correlation measures.

#### Psychological mediation analyses

We found a significant positive association between BMI and the BIS subscales (BIS total: β=0.18, CI: [0.06; 0.28], p=0.0016; BIS motor: β=0.06, CI: [0.013; 0.10], p=0.0013 and BIS self-control: β=0.06, CI: [0.017; 0.10], p=0.0046). Yet, none of the scales significantly mediated the relationship between BMI and mFD (see Table 2 and Figure 3),

**Table 2:**
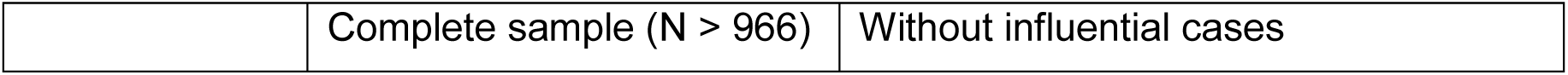
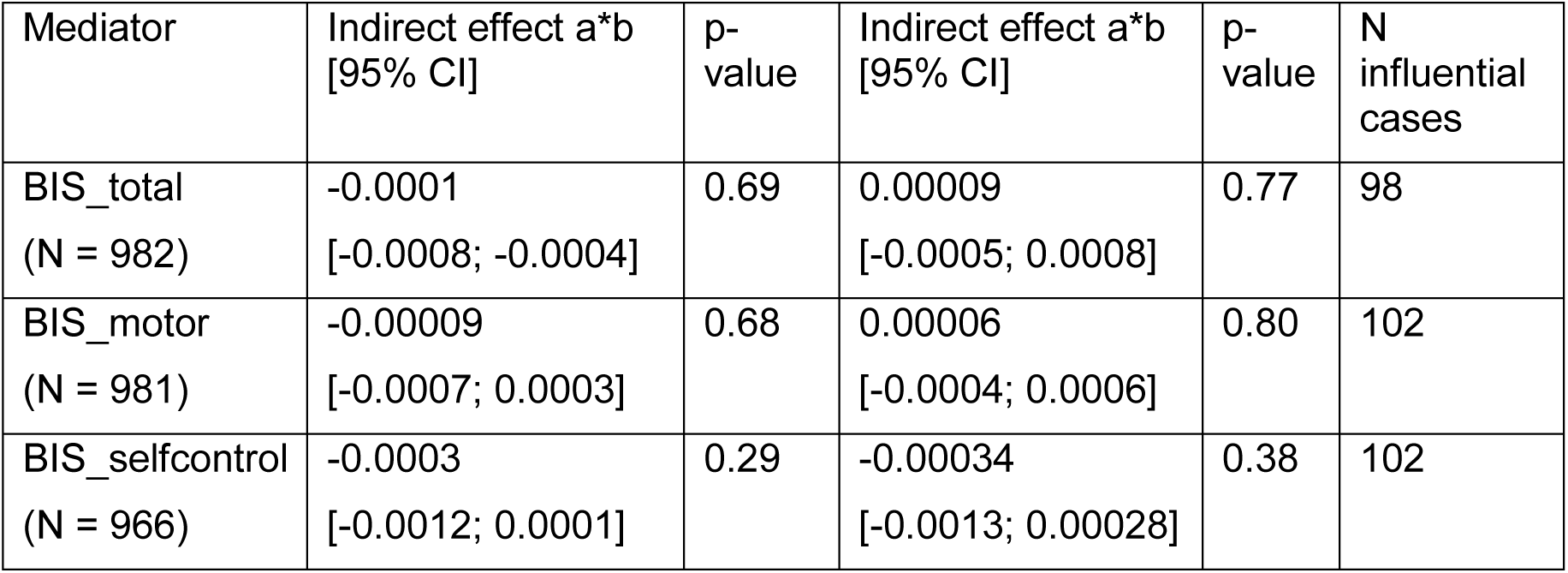
Results of the psychological mediation models, shown for the full samples (sample size depending on investigated variable, see left column) and without influential cases. BIS: Barratt Impulsiveness Scale; CI: confidence interval

| Mediator | Indirect effect a*b<br>[95% CI] | p-value | Indirect effect a*b<br>[95% CI] | p-value | N<br>influential<br>cases |
| --- | --- | --- | --- | --- | --- |
| BIS_total<br>(N = 982) | -0.0001<br>[-0.0008; -0.0004] | 0.69 | 0.00009<br>[-0.0005; 0.0008] | 0.77 | 98 |
| BIS_motor<br>(N = 981) | -0.00009<br>[-0.0007; 0.0003] | 0.68 | 0.00006<br>[-0.0004; 0.0006] | 0.80 | 102 |
| BIS_selfcontrol<br>(N = 966) | -0.0003<br>[-0.0012; 0.0001] | 0.29 | -0.00034<br>[-0.0013; 0.00028] | 0.38 | 102 |

**Figure 3:**
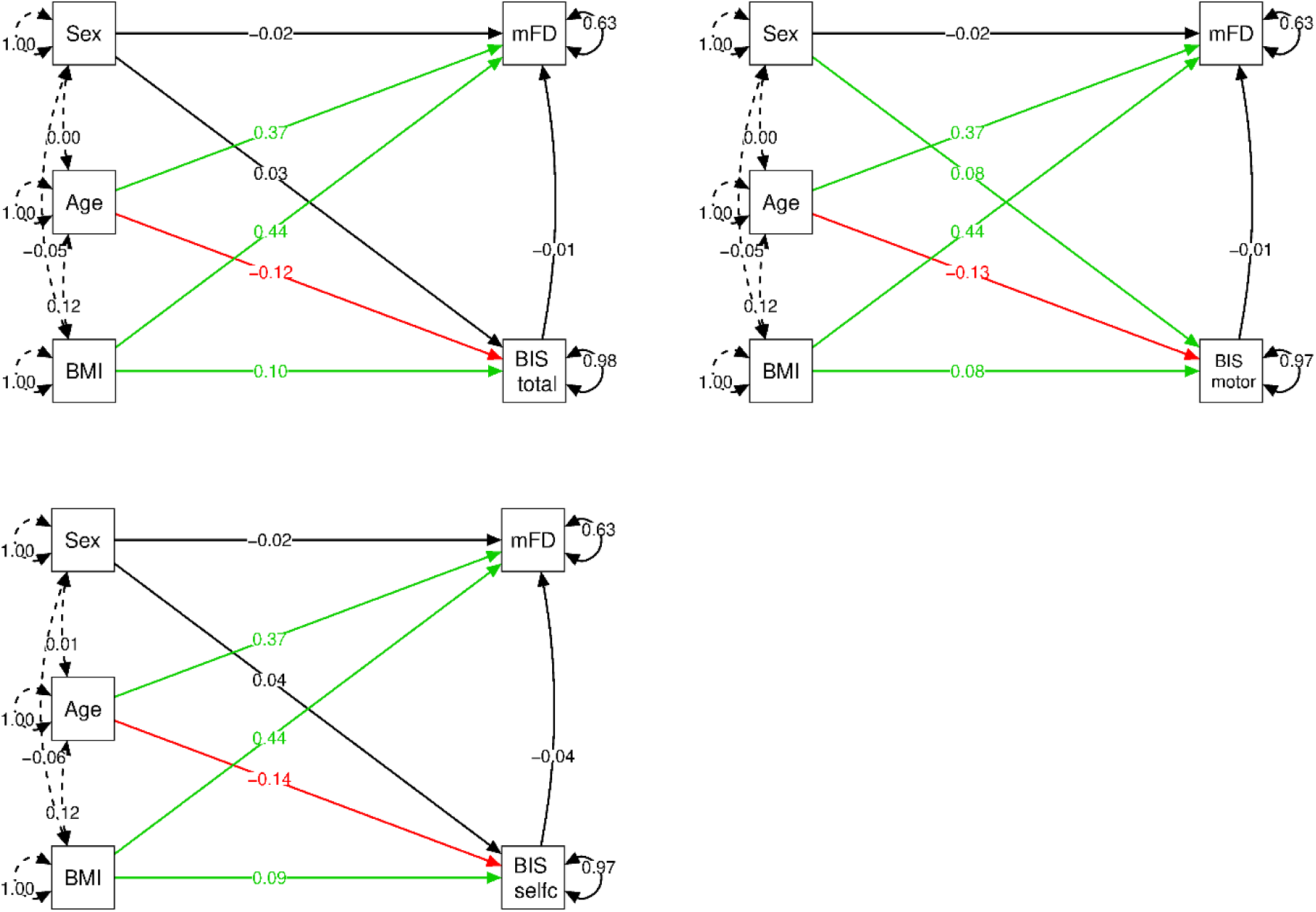
Mediation model for BIS total, motor and self-control impulsivity scales showing standardized estimates. Dashed arrows indicate residual (co-)variance. Solid arrows indicate estimated regression coefficients, green/red indicate significant positive/negative associations (p<0.05), black indicates no significant association. The reference category for sex was male. BMI: body mass index, mFD: mean framewise displacement, BIS: Barratt Impulsiveness Scale.

## 2.4 Discussion

In this meta-GWAS including > 35,000 individuals, we did not find a statistically significant genetic variant associated with mean or maximal FD. While there was a positive correlation of BMI and head motion in a subsample analysis of ∼1000 individuals, neither physiological nor psychological measures mediated this association. These results do therefore not support that single factors underlie the association of BMI and head motion. The phenotypic association of BMI and head motion could be rather related to more complex factors, e.g. polygenetic traits with many low-effect variants or different sources of head motion, i.e. specific breathing patterns.

In the meta-GWAS of head motion, we expected to detect variants related to obesity (e.g. FTO) or impulsivity (e.g. ADHD-associated SNPs ADGRL3). While one of the sub-threshold variants is also weakly linked with body weight according to the UK Biobank gene atlas, no single variant is reliably associated with head motion in this well-powered analysis. This indicates that the genetic correlation of head motion and BMI/impulsivity which has been previously reported is driven by polygenic effects with very small individual effect sizes (Couvy-Duchesne et al., 2016; Hodgson et al., 2016). Importantly, the previous studies estimated heritability of head motion from family structure, not genome-wide significant SNPs. For polygenetic traits, GWAS-based heritability is smaller than family-based heritability due to large number of small effect SNPs, untagged variants with larger effects sizes, non-additive genetic variation and environmental effects (including epigenetic modifications) (Yang et al., 2017). Thus, it is likely that no single variant explains enough variance to be a genome-wide predictor of this polygenic trait. Even though head motion was determined similarly across studies, it is possible that study-specific patient positioning and fixation influenced the magnitude of head motion and inter-participant variation, and with that, decreased our power due to increased heterogeneity between studies. In the UKBB, the largest sample in our analysis, the correlation of BMI and mean FD was comparable to previous reports (Pearson’s r ∼ 0.5, data not shown). Thus, differences in data acquisition are unlikely to explain the null result.

Previous studies indicated that physiological rather than psychological factors are major determinants of head motion (Ekhtiari et al., 2019; Makowski et al., 2019). However, in our pre-registered analyses, neither physiological nor psychological measures independently mediated the link between BMI and head motion.

In line with the literature, we found BMI to be associated with higher trait impulsivity in our sample (Meule & Blechert, 2016). Yet, impulsivity was not related to head motion in our sample. Possibly, the link between impulsivity and head motion is stronger in children and young adults who may have less self-control and for whom other (physiological) factors do not yet play an important role (Couvy-Duchesne et al., 2016; Kong et al., 2014).

In the physiological mediation models, there was no mediating effect of respiratory rate, volume per time and heart rate. This might be due to two aspects. First, the hypothesized relationship between BMI and respiratory parameters was largely absent. In the literature, alterations in the respiratory system have been mostly reported for morbidly obese individuals (BMI > 40 kg/m^2^) (Littleton, 2012). In our sample of 1006 individuals, there were only 8 morbidly obese individuals and 40 individuals with a BMI above 35 kg/m^2^. Thus, although we saw a trend association of BMI and RR, differences in respiration in our sample may not have been as pronounced.

Second, we expected higher HR and RR to predict higher head motion, but found no or even the inverse relationship. Upon inspection of exemplary carpet plots, we saw that participants with lower RR often showed irregular breathing patterns with many deep breaths which were associated with higher head motion. Participants with higher RR had more regular respiration cycles, which went along with lower head motion (see Figure 4)

**Figure 4:**
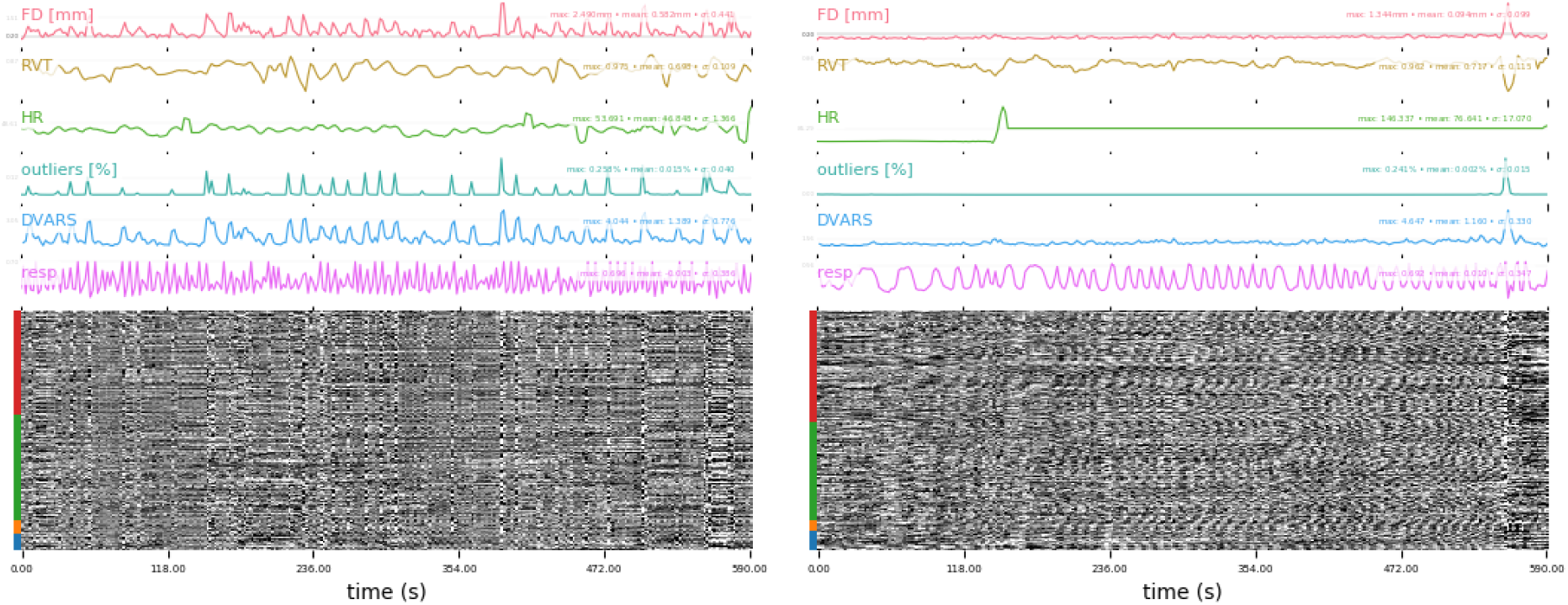
Motion (FD, outliers %), physiological (RVT, respiration trace and HR) and fMRI signal (DVARS) parameters shown over individual voxel timeseries (red: GM, green: WM, yellow: ventricle, blue: cerebellum). For the left participant, RR is 0.18 and average mFD is 0.29 mm, for the right participant RR is 0.41 and average mFD is 0.004).

Interestingly, we found higher BMI to be associated with less correlation of frame-to-frame head motion and RVT. At average BMI, this correlation was close to zero, indicating that FD and RVT become less associated with higher BMI. This might indicate that FD depends on factors other than respiration with higher BMI, or that episodes with respiration-related head motion and subsequent deep breathing phase with decreased RVT (in which anti-correlation of FD and RVT have been observed) are more frequent in individuals with higher BMI. A major strength of this study is the large meta-GWAS of a rsfMRI-derived phenotype in 7 cohorts. Furthermore, we conducted a comprehensive, pre-registered investigation of physiological and psychological mediators of the association between BMI and head motion. The most important limitations of the meta-GWAS study are the biased sample sizes among the included studies and the absence of a replication sample while for the mediation analyses a more detailed investigation of breathing patterns, more specific psychological measures and the inclusion of a proxy for anxiety in the scanner would have been desirable. We conclude that the strong phenotypic correlation of head motion and BMI is not induced by strong genetic variants, nor explained by simple physiological or psychological variables on a population level. Further studies might explore specific breathing patterns associated with head motion, and investigate further potential sources of the strong phenotypic link between BMI and head motion. In the meantime, rsfMRI studies investigating BMI or aging should prevent head motion during acquisition using customized head molds or prospective motion correction and employ adequate motion correction techniques during preprocessing to avoid a confounding effect of head motion on their neuroimaging measures.

## References

Beyer, F., Kharabian Masouleh, S., Huntenburg, J. M., et al. (2017). Higher body mass index is associated with reduced posterior default mode connectivity in older adults. Hum Brain Mapp, 38(7), 3502–3515. doi:10.1002/hbm.23605

Birn, R. M., Diamond, J. B., Smith, M. A., & Bandettini, P. A. (2006). Separating respiratory-variation-related fluctuations from neuronal-activity-related fluctuations in fMRI. Neuroimage, 31(4), 1536–1548.

Burki, N. K., & Baker, R. W. (1984). Ventilatory regulation in eucapnic morbid obesity. The American review of respiratory disease, 129(4), 538–543.

Chang, C., Cunningham, J. P., & Glover, G. H. (2009). Influence of heart rate on the BOLD signal: the cardiac response function. Neuroimage, 44(3), 857–869.

Couvy-Duchesne, B., Blokland, G. A. M., Hickie, I. B., et al. (2014). Heritability of head motion during resting state functional MRI in 462 healthy twins. Neuroimage, 102, 424–434. doi:https://doi.org/10.1016/j.neuroimage.2014.08.010

Couvy-Duchesne, B., Ebejer, J. L., Gillespie, N. A., et al. (2016). Head Motion and Inattention/Hyperactivity Share Common Genetic Influences: Implications for fMRI Studies of ADHD. PLoS One, 11(1), e0146271. doi:10.1371/journal.pone.0146271

Ekhtiari, H., Kuplicki, R., Yeh, H.-w., & Paulus, M. P. (2019). Physical characteristics not psychological state or trait characteristics predict motion during resting state fMRI. Sci. Rep., 9(1), 419. doi:10.1038/s41598-018-36699-0

Engelhardt, L. E., Roe, M. A., Juranek, J., et al. (2017). Children’s head motion during fMRI tasks is heritable and stable over time. Dev. Cogn. Neurosci., 25, 58–68. doi:https://doi.org/10.1016/j.dcn.2017.01.011

Fair, D. A., Miranda-Dominguez, O., Snyder, A. Z., et al. (2020). Correction of respiratory artifacts in MRI head motion estimates. Neuroimage, 208, 116400. doi:https://doi.org/10.1016/j.neuroimage.2019.116400

Heidari, S., Babor, T. F., De Castro, P., Tort, S., & Curno, M. (2016). Sex and gender equity in research: rationale for the SAGER guidelines and recommended use. Research integrity and peer review, 1(1), 1–9.

Hodgson, K., Poldrack, R. A., Curran, J. E., et al. (2016). Shared Genetic Factors Influence Head Motion During MRI and Body Mass Index. Cereb. Cortex.

Kasper, L., Bollmann, S., Diaconescu, A. O., et al. (2017). The PhysIO toolbox for modeling physiological noise in fMRI data. J. Neurosci. Methods, 276, 56–72.

Kong, X. Z., Zhen, Z., Li, X., et al. (2014). Individual differences in impulsivity predict head motion during magnetic resonance imaging. PLoS One, 9(8). doi:10.1371/journal.pone.0104989

Littleton, S. W. (2012). Impact of obesity on respiratory function. Respirology, 17(1), 43–49.

Loeffler, M., Engel, C., Ahnert, P., et al. (2015). The LIFE-Adult-Study: objectives and design of a population-based cohort study with 10,000 deeply phenotyped adults in Germany. BMC Public Health, 15(1), 691. doi:10.1186/s12889-015-1983-z

Madan, C. R. (2018). Age differences in head motion and estimates of cortical morphology. PeerJ, 6(2167-8359 (Print)), e5176. doi:10.7717/peerj.5176

Makowski, C., Lepage, M., & Evans, A. C. (2019). Head motion: the dirty little secret of neuroimaging in psychiatry. Journal of psychiatry & neuroscience: JPN, 44(1), 62.

Meule, A., & Blechert, J. (2016). Trait impulsivity and body mass index: a cross-sectional investigation in 3073 individuals reveals positive, but very small relationships. Health Psychology Open, 3(2), 2055102916659164.

Patton, J. H., Stanford, M. S., & Barratt, E. S. (1995). Factor structure of the Barratt impulsiveness scale. J. Clin. Psychol., 51(6), 768–774. doi:10.1002/1097-4679(199511)51:6<768::aid-jclp2270510607>3.0.co;2-1

Power, J., Schlaggar, B., & Petersen, S. (2015). Recent progress and outstanding issues in motion correction in resting state fMRI. Neuroimage, 105, 536–551.

Power, J. D., Barnes, K. A., Snyder, A. Z., Schlaggar, B. L., & Petersen, S. E. (2012). Spurious but systematic correlations in functional connectivity MRI networks arise from subject motion. Neuroimage, 59(3), 2142–2154. doi:10.1016/j.neuroimage.2011.10.018

Savalia, N. K., Agres, P. F., Chan, M. Y., et al. (2017). Motion-related artifacts in structural brain images revealed with independent estimates of in-scanner head motion. Human brain mapping, 38(1), 472–492.

Siegel, J. S., Mitra, A., Laumann, T. O., et al. (2017). Data Quality Influences Observed Links Between Functional Connectivity and Behavior. Cereb. Cortex, 27(9), 4492–4502. doi:10.1093/cercor/bhw253

Thomson, P., Johnson, K. A., Malpas, C. B., et al. (2020). Head Motion During MRI Predicted by out-of-Scanner Sustained Attention Performance in Attention-Deficit/Hyperactivity Disorder. Journal of Attention Disorders, 1087054720911988. doi:10.1177/1087054720911988

Vainik, U., Baker, T. B., Dadar, M., et al. (2018). Neurobehavioural Correlates of Obesity are Largely Heritable. bioRxiv, 204917.

Van Dijk, K. R., Sabuncu, M. R., & Buckner, R. L. (2012). The influence of head motion on intrinsic functional connectivity MRI. Neuroimage, 59(1), 431–438.

van Minde, D., Klaming, L., & Weda, H. (2014). Pinpointing moments of high anxiety during an MRI examination. Int. J. Behav. Med., 21(3), 487–495.

Yang, J., Zeng, J., Goddard, M. E., Wray, N. R., & Visscher, P. M. (2017). Concepts, estimation and interpretation of SNP-based heritability. Nat. Genet., 49(9), 1304–1310. doi:10.1038/ng.3941

Zeng, L. L., Wang, D., Fox, M. D., et al. (2014). Neurobiological basis of head motion in brain imaging. Proc Natl Acad Sci USA, 111(16), 6058–6062. doi:10.1073/pnas.1317424111

