## Supplementary Material for "Investigating the genetic and environmental basis of head micromovements during MRI"

**Supplementary Table 1:** Information on population characteristics, MRI acquisition and head motion preprocessing in the participating studies

| Study | N (total/female/male) | Age (mean (sd) in years) | Mean framewise displacement (mean (sd) in mm) | Scanner (field strength) | Sequence (TR, parallel acquisition, multiband) | Motion correction tool |
|---|---|---|---|---|---|---|
| LIFE-Adult Baseline | 2083 (1022/1061) | 57.66 (15.3) | 0.33 (0.16) | Siemens Verio, 3 T | BOLD-EPI, (2s, 1, none ) | MCFLIRT |
| Rotterdam Study 1 | 372 (207/165) | 82.4 (3.6) | 1.19 (1.85) | Signa Excite, 1.5 T | BOLD-EPI, (2.9s, 1, none ) | MCFLIRT |
| Rotterdam Study 2 | 270 (131/139) | 73.4 (4.5) | 0.97 (1.34) | Signa Excite, 1.5 T | BOLD-EPI, (2.9s, 1, none ) | MCFLIRT |
| Rotterdam Study 3 | 1743 (946/797) | 62.0 (5.8) | 0.80 (0.96) | Signa Excite, 1.5 T | BOLD-EPI, (2.9s, 1, none ) | MCFLIRT |
| ASPS-Fam | 164 (97/67) | 67.3 (10.6) | 0.17 (0.11) | Siemens Trio Tim, 3T | BOLD-EPI, (3s, 1, none ) | MCFLIRT |
| SHIP | 765(410/355) | 58.1 (12.1) | 0.18 (0.09) | Siemens Avanto, 1.5 T | BOLD-EPI, (2.9 s, 1, none) | 3dvolreg |
| UKBB | 29,778(15728/13996) | 63.8 (7.5) | 0.20 (0.09) | Siemens Skyra 3T | BOLD-EPI (0.735s, 8, none) | MCFLIRT |

**Supplementary Table 2:** Information on genotyping and genetic analysis (HWE: Hardy-Weinberg Equilibrium, MAF: minor allele frequency, SNP: single nucleotid polymorphism)

| **Cohort** | **Analysis** | **HWE** | **MAF** | **SNP call rate** | **Imputation method** | **Reference panel** | **Genotype Platform** |
|---|---|---|---|---|---|---|---|
| LIFE-Adult | Snptest version 2.5.2 | < 10−6 | ≥0.02 | >97% | impute2, version 2.3.2 | 1000 Genomes (phase 3v5) | Affymetrix Axiom Genome-Wide CEU 1 |
| SHIP | QUICKTEST version 0.9 | < 0.001 | >0.01 | >0.8 | IMPUTE v2.2.2 | 1000 Genomes (phase 1v3) | Affymetrix Human SNP Array 6.0 |
| Rotterdam Study I | ProbABEL v.0.4.4 | < 10−6 | > 0.01 | > 0.98 | MaCH (Mach 1.0.18.c) and minimac (2012.8.15) | 1000G Phase1v3 all ethnicities | Illumina HumanHap550K(+Duo) and 610-Quad BeadChip |
| Rotterdam Study II | ProbABEL v.0.4.4 | < 10−6 | > 0.01 | > 0.98 | MaCH (Mach 1.0.18.c) and minimac (2012.8.15) | 1000G Phase1v3 all ethnicities | Illumina HumanHap550-Duo BeadChip |
| Rotterdam Study III | ProbABEL v.0.4.4 | < 10−6 | > 0.01 | > 0.98 | MaCH (Mach 1.0.18.c) and minimac (2012.8.15) | 1000G Phase1v3 all ethnicities | Illumina HumanHap610K-Quad |
| ASPS-Fam | ProbABEL | < 5*10-6 | > 0.05 | > 98% | IMPUTE2 v2.3.1 | 1000G ALL phase1 integrated variant set realease v3 (march 2012) | Affymetrix Genome-Wide Human SNP Array 6.0 |
| UKBB | BOLT-LMM 2.3.4 | none by analyst (UV) | BOLT filtered 39458 SNPs with > 0.1 missing, and 744798 called | >0.97% | IMPUTE4 | Haplotype Reference Consortium (HRC) and UK10K haplotype resource | Applied Biosystems UK Biobank Axiom Array |

| | | | |
|---|---|---|---|
| | | | SNPs remaining |

Supplementary Figure 1: Strongest effects for mean FD in rs2218592 on chromosome 3.

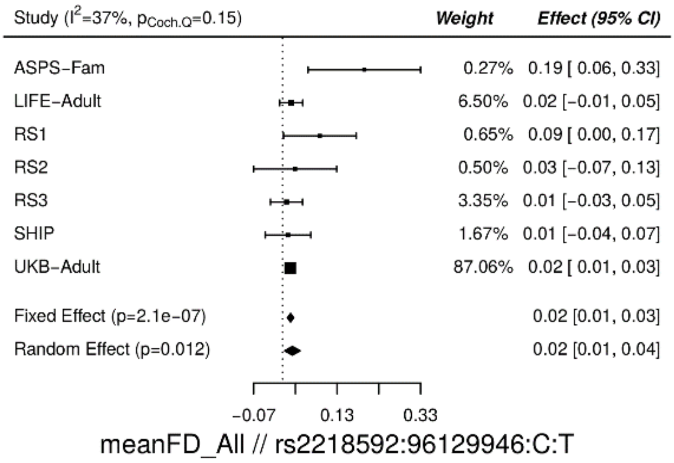

Supplementary Figure 2: Second strongest effects for mean FD in rs8054401 on chromosome 16.

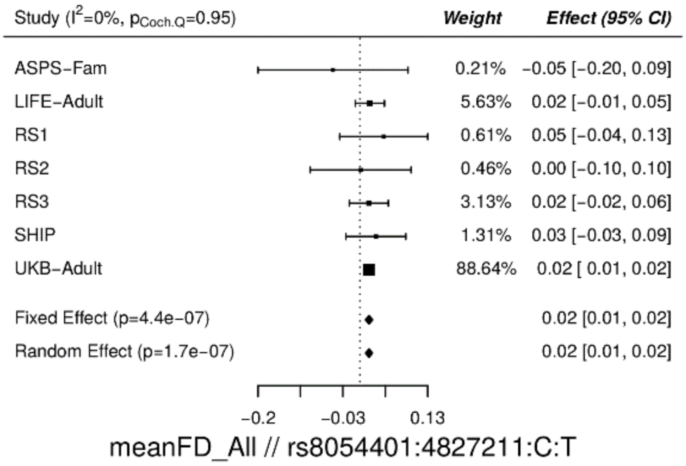

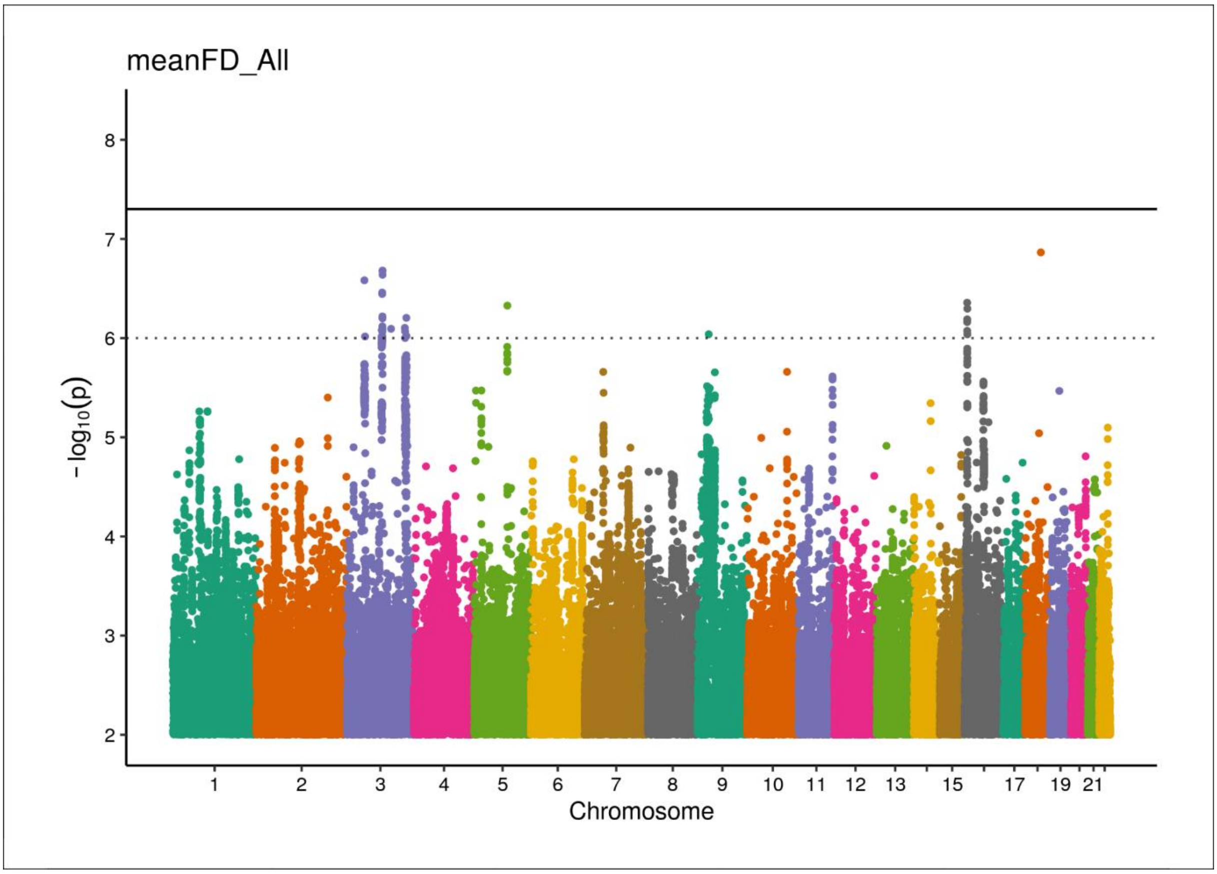

Supplementary Figure 3: Manhattan plot from the meta-GWAS for mean FD phenotype across males and females.

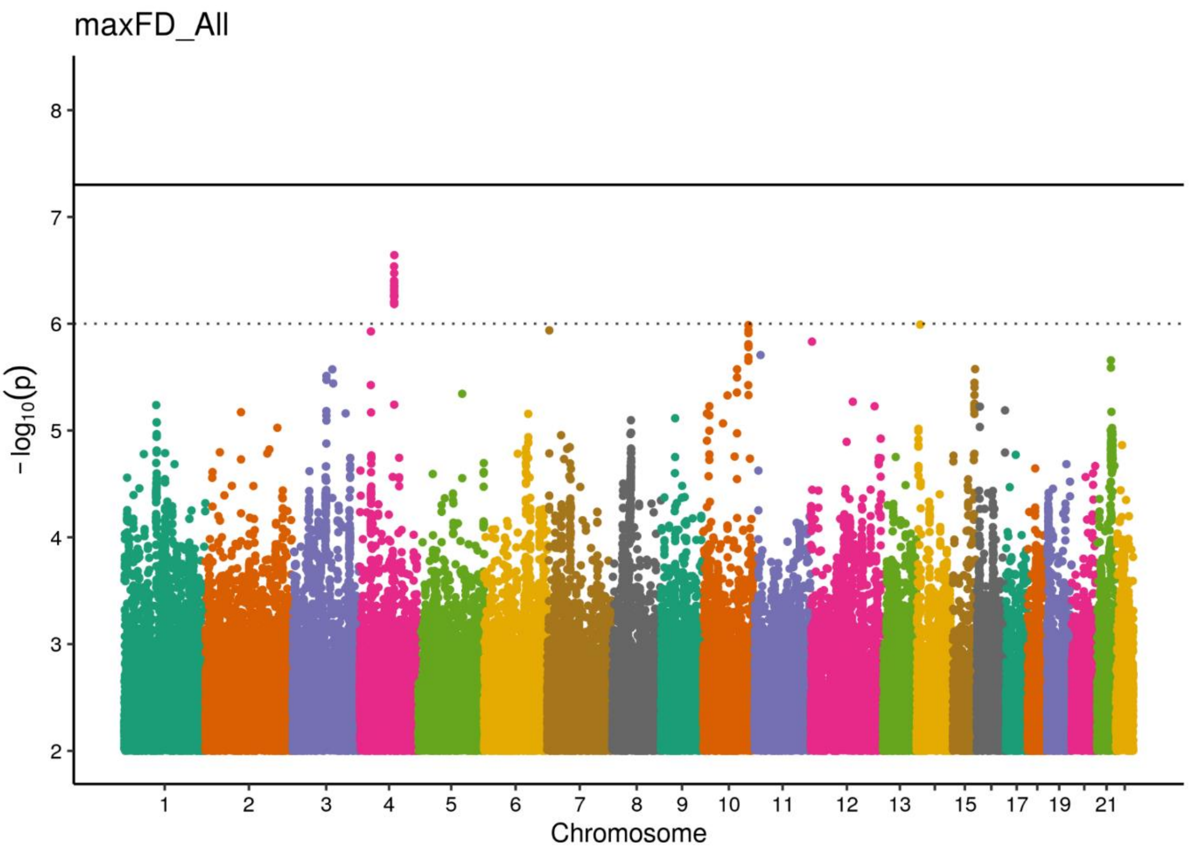

Supplementary Figure 4: Manhattan plot from the meta-GWAS for maximal FD phenotype across males and females.

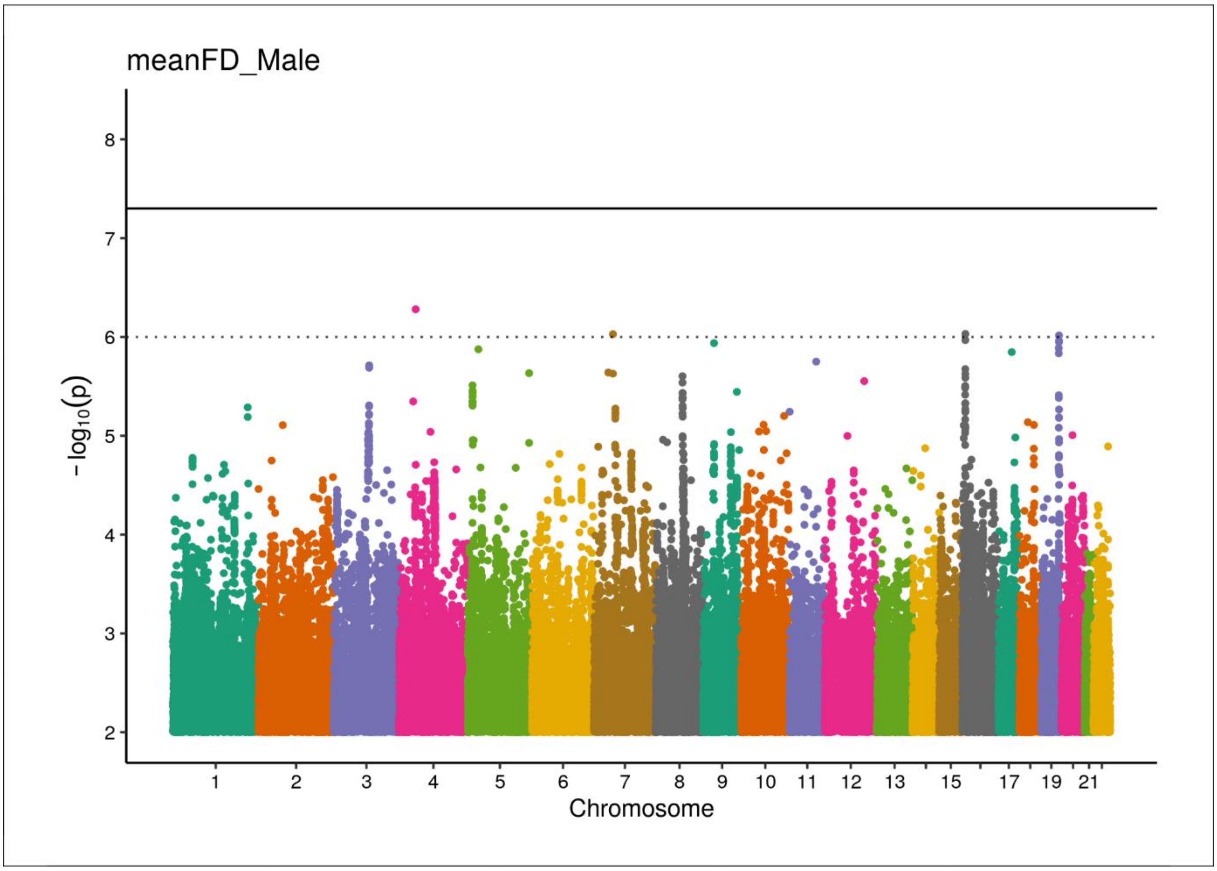

Supplementary Figure 5: Manhattan plot from the meta-GWAS for mean FD phenotype in men.

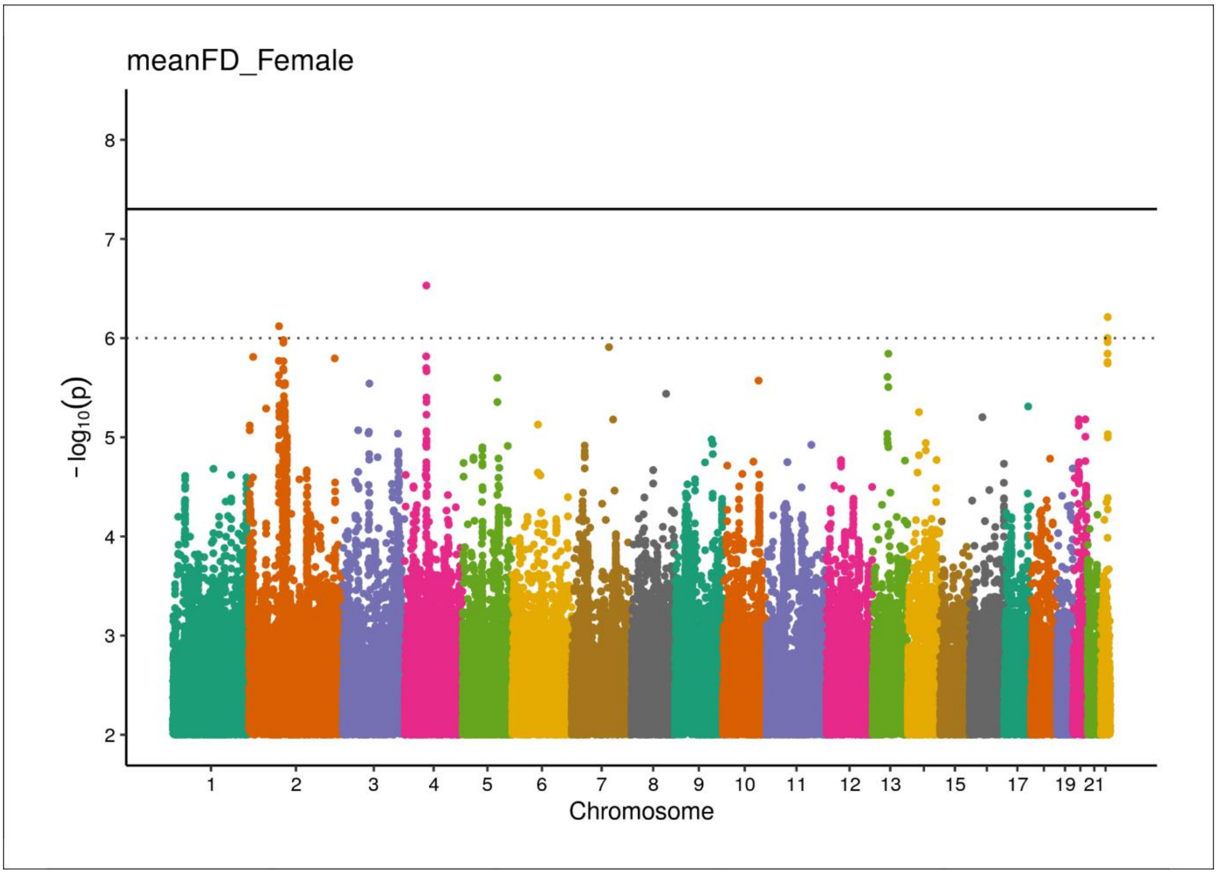

Supplementary Figure 6: Manhattan plot from the meta-GWAS for mean FD phenotype in women.

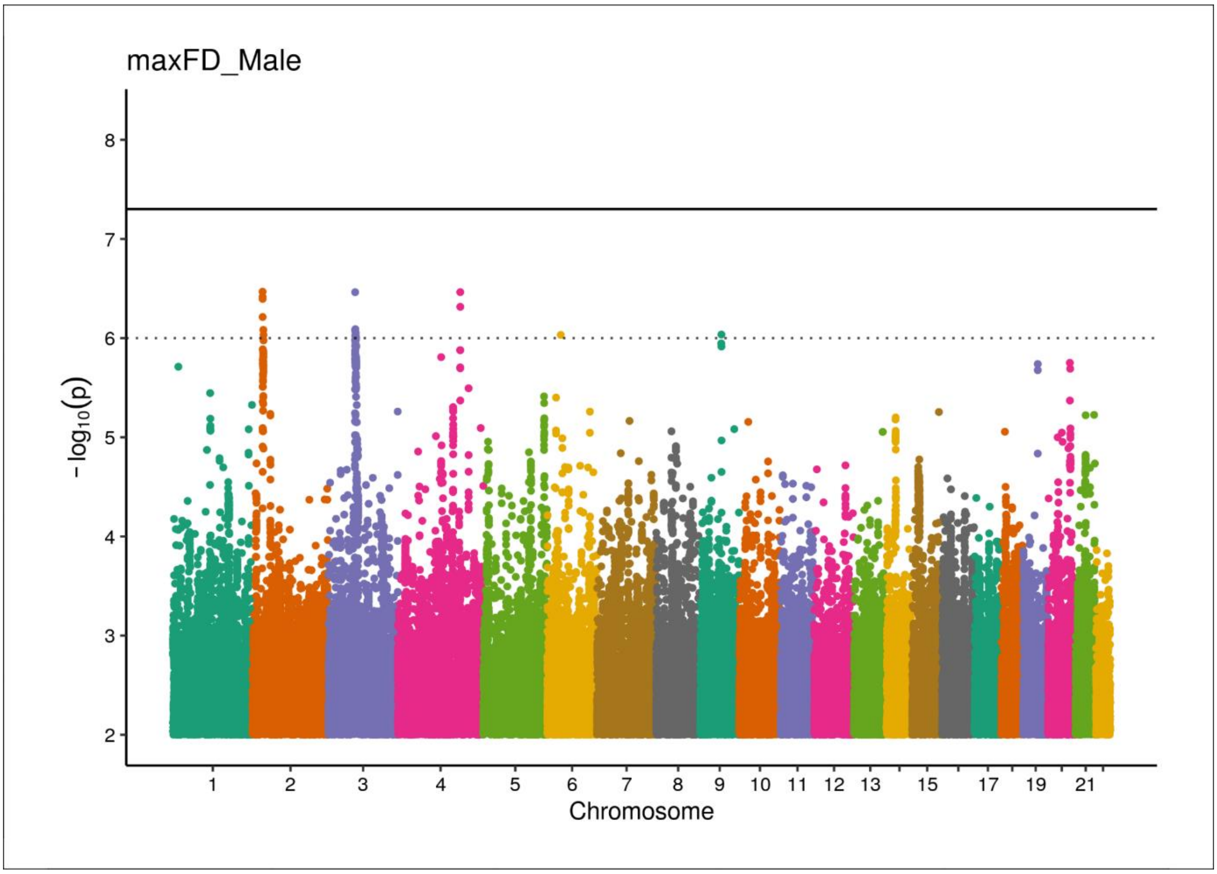

Supplementary Figure 7: Manhattan plot from the meta-GWAS for maximal FD phenotype in men.

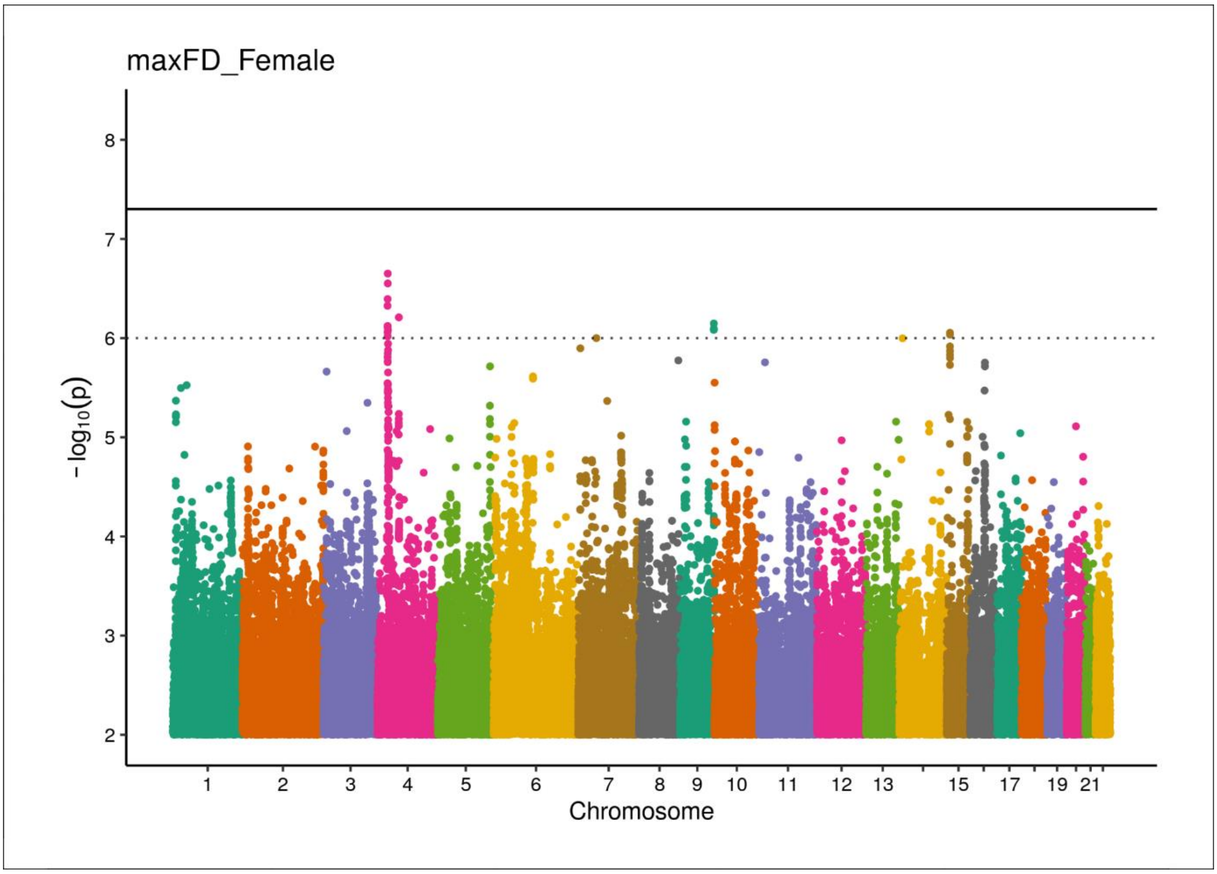

Supplementary Figure 8: Manhattan plot from the meta-GWAS for maximal FD phenotype in women.

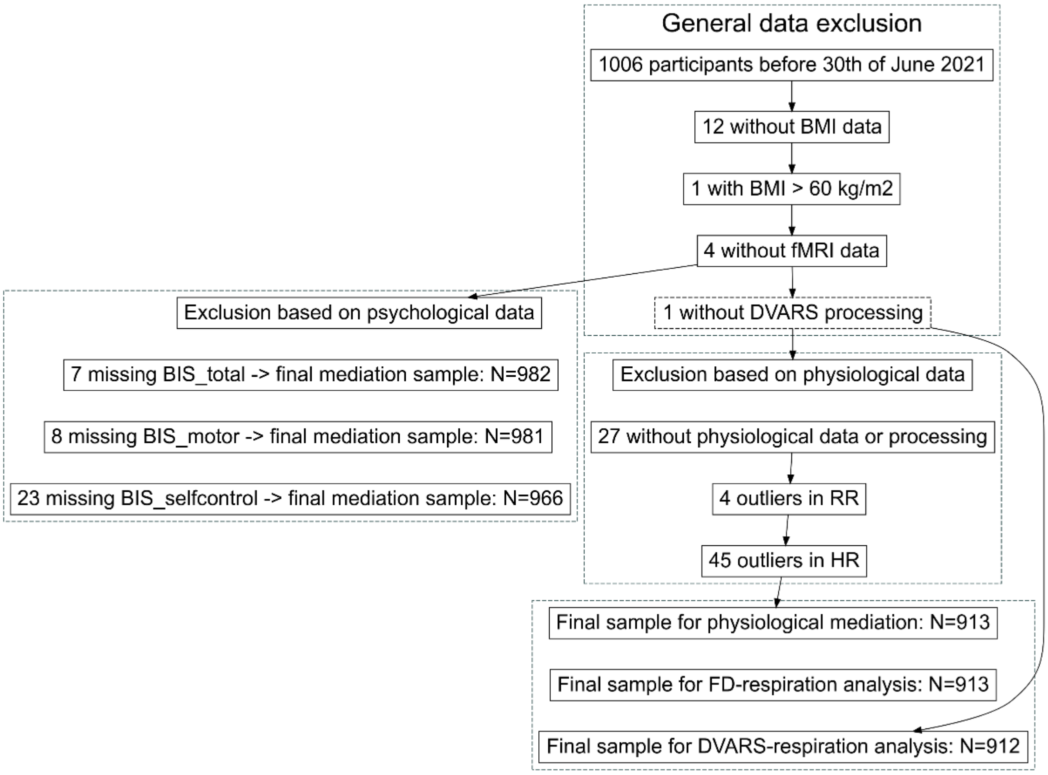

Supplementary Figure 9: Flowchart showing the sample sizes for physiological and psychological mediation and regression analyses in the LIFE-Adult study.
